# PRMT5 is required for full-length *HTT* expression by repressing multiple proximal intronic polyadenylation sites

**DOI:** 10.1101/2024.03.14.584861

**Authors:** Mona A. AlQazzaz, Felipe E. Ciamponi, Jolene C. Ho, Maxim I. Maron, Manisha Yadav, Aiden M. Sababi, Graham MacLeod, Moloud Ahmadi, Garrett Bullivant, Vincent Tano, Sarah R. Langley, María Sánchez-Osuna, Patty Sachamitr, Michelle Kushida, Laura Richards, Costanza Ferrari Bardile, Mahmoud A. Pouladi, Trevor Pugh, Mike Tyers, Stephane Angers, Peter B. Dirks, Gary D. Bader, Katlin B. Massirer, Dalia Barsyte-Lovejoy, David Shechter, Rachel J. Harding, Cheryl H. Arrowsmith, Panagiotis Prinos

## Abstract

Expansion of the CAG trinucleotide repeat tract in exon 1 of the *Huntingtin* (*HTT*) gene above a threshold of ∼36 repeats causes Huntington’s disease (HD) through the expression of a polyglutamine-expanded form of the HTT protein. This mutation triggers wide-ranging cellular and biochemical pathologies leading to cognitive, motor, and psychiatric symptoms in HD patients. As accurate splicing is required to produce the full-length HTT protein of ∼348 kDa, targeting *HTT* splicing with small molecule drugs is a compelling approach to lower HTT protein levels to treat HD, and splice modulators are being tested in the clinic. Here, we identify PRMT5 as a novel regulator of *HTT* mRNA splicing and alternative polyadenylation. PRMT5 inhibition disrupts the splicing of *HTT* introns 9 and 10, leading to activation of multiple proximal intronic polyadenylation sites within these introns and promoting premature termination, cleavage and polyadenylation (PCPA) of the *HTT* mRNA, thus lowering total HTT protein levels. We also detected increasing levels of these truncated, intron-containing *HTT* transcripts across a series of neuronal differentiation samples which correlated with lower PRMT5 expression. Notably, PRMT5 inhibition in glioblastoma (GBM) stem cells potently induced neuronal differentiation. We posit that PRMT5-mediated regulation of intronic polyadenylation, premature termination and cleavage of the *HTT* mRNA modulates HTT expression and plays an important role during embryonic development and neuronal differentiation.

## Introduction

Huntington’s disease (HD) is an autosomal dominant inherited neurodegenerative disease displaying a range of psychiatric, cognitive and physical symptoms^1^ that affects ∼1 in 10,000 people^2^. Characterised by progressive loss of striatal neurons^3^, HD is caused by expansion of the CAG trinucleotide repeat tract in exon 1 of the *Huntingtin* (*HTT*) gene above a threshold of ∼36 repeats, leading to the expression of a polyglutamine-expanded form of the HTT protein. Deletion of *Htt* is embryonically lethal in mice^5^ and the protein is known to play important roles in numerous cellular processes that are critical for neuronal development^6^. The *HTT* locus spans 167 kb, encompassing 67 exons and 66 introns; thus, accurate RNA splicing regulation is essential to produce the mature and functional HTT protein. In an HD context, the expression of an expanded and truncated *HTT* intron 1 (*HTT1a*) by aberrant splicing leads to the accumulation of a short polyadenylated mRNA which is translated to produce the toxic exon 1 protein fragment^7^. Other groups have reported a variety of *HTT* isoforms produced by alternative splicing ^8–14^. More recently, splicing modulation has emerged as a promising therapeutic approach for lowering levels of HTT, a strategy being tested in clinical trials to slow or halt HD progression^15,16^. The small molecule splicing modulators, Branaplam and PTC-518, promote inclusion of a poison pseudoexon within intron 49 in the *HTT* transcript, leading to the introduction of premature termination codons and nonsense-mediated decay (NMD) of the *HTT* mRNA, ultimately resulting in the lowering of HTT protein levels^17–19^. Nevertheless, the endogenous factors that regulate *HTT* splicing in normal physiology and development remain largely unknown.

Protein arginine methyltransferase 5 (PRMT5) catalyses the symmetric dimethylation of arginine residues on target proteins including histones, RNA-binding proteins, splicing factors, transcription factors and their complexes, DNA damage and repair factors, and metabolic regulators. Major PRMT5 cellular substrates include the spliceosomal Sm proteins SmD1 (SNRPD1), SmD3 (SNRPD3), and SmB/Bʹ (SNRPB)^22–24^. Symmetric arginine methylation of these snRNP proteins is required for their efficient binding to the SMN protein and assembly into the mature spliceosome^23^. We and others have demonstrated the importance of PRMT5 for the maintenance of RNA splicing fidelity^25–29^. PRMT5 inhibition results in widespread intron retention leading to global downregulation of proteins implicated in cell proliferation^26,28^. Interestingly, both PRMT5 and HTT are essential for neuronal stem cell proliferation and self-renewal^25,30,>3113,18,19^. Since splicing and RNA metabolism have been shown to play a central role in neurogenesis, neuronal differentiation, and neurodegenerative diseases^33–38^ it is plausible that PRMT5 may regulate these processes via RNA splicing, and that this role may intersect with HTT.

We had previously identified PRMT5 as a regulator of splicing and stemness in glioblastoma (GBM) stem cells and identified hundreds of intron retention events as the most prominent effect of PRMT5 inhibition on splicing^28^. Global proteome surveying linked the mis-splicing induced by PRMT5 inhibition to a global lowering of proteins involved in cell proliferation. Inspired by this and the recent discoveries of splicing inhibitors as HTT-lowering agents, we investigated the effects of PRMT5 inhibition on *HTT* mRNA splicing. Here, we show that PRMT5 inhibition impairs splicing of *HTT* introns 9 and 10, leading to the activation of a cluster of cryptic alternative polyadenylation sites within these introns, resulting in premature cleavage and polyadenylation (PCPA) of the *HTT* mRNA and reduction of total HTT protein levels. Strikingly, we found increased levels of this truncated HTT isoform during neuronal differentiation concomitant with lower PRMT5 levels, suggesting a physiological role for this regulation.

## Results

### PRMT5 inhibition decreases *HTT* mRNA and protein levels in GBM models

We previously reported that PRMT5 inhibition impaired splicing and stemness in a large panel of patient-derived GBM stem cells. RNA sequencing of GBM stem cells treated with 1 μM GSK591, a potent and selective PRMT5 inhibitor, revealed widespread mis-splicing, with intron retention events being the most prevalent splicing alteration observed. One of the top affected genes in this study was *HTT* (**Figure 1a**), which was significantly down-regulated following PRMT5 inhibition with GSK591 in these lines. Proteomics analysis in three different patient-derived GBM cell lines (G561, G564 and G583) also showed decreased HTT protein with PRMT5 inhibition with 1 μM GSK591 or LLY-283 (**Figure 1a**). Building upon this finding, we validated *HTT* mRNA downregulation by quantitative reverse transcription PCR (RT-qPCR, referred herein as qPCR) in three different GBM stem cell lines (**Figure 1b**). The levels of *HTT* mRNA were significantly lower after treatment of GBM cells with either 1 µM GSK591 or LLY-283 (**Figure 1b**), both PRMT5 inhibitors, but not with an inactive control compound, SGC2096. Western blots of cell lysates from two patient-derived GBM lines treated with these compounds showed consistent decrease in HTT protein following PRMT5 inhibition, thus corroborating the mRNA decrease with either GSK591 or LLY-283 (**Figure 1c, d**).

**Figure 1.**
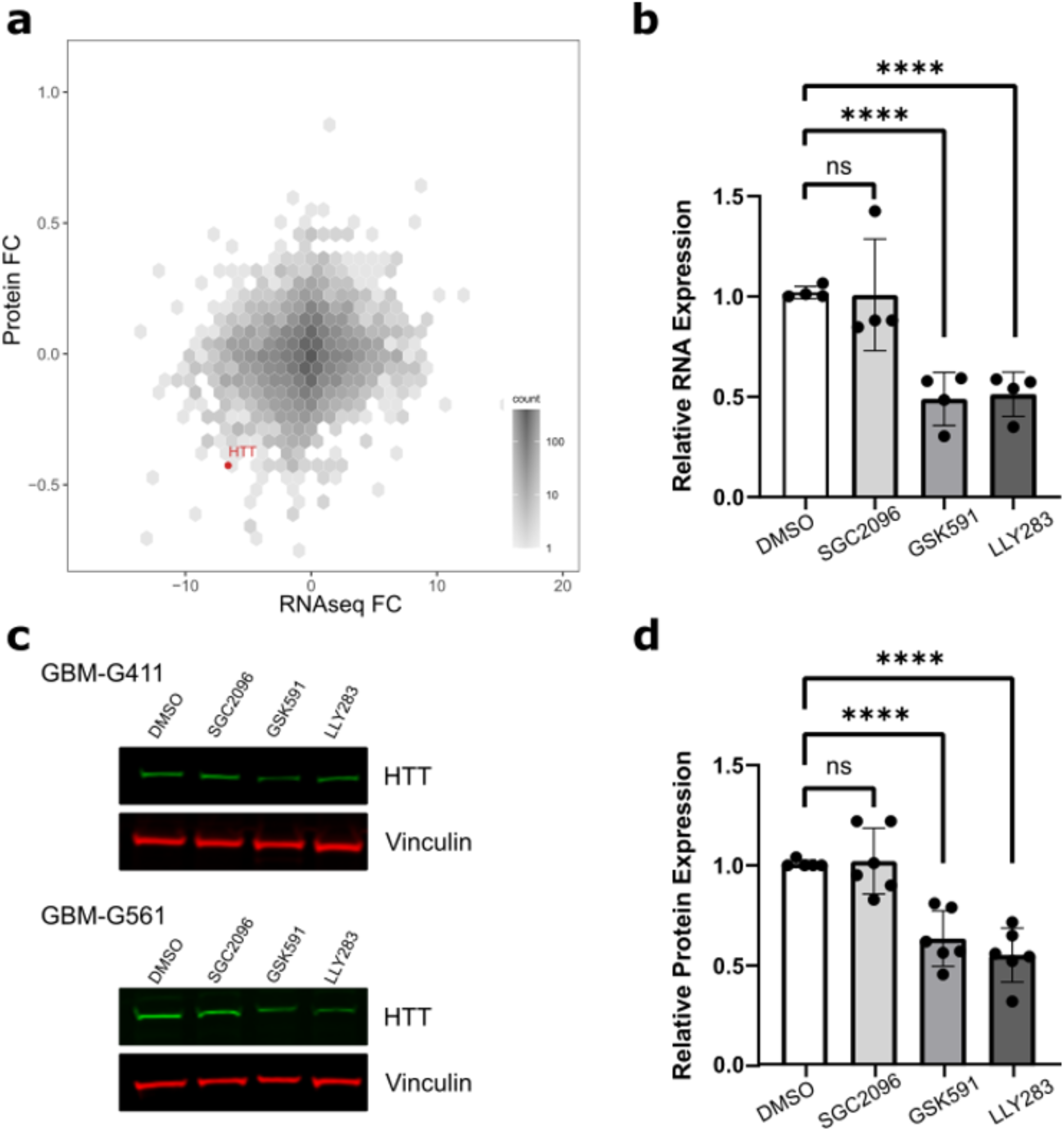
PRMT5 inhibitor treatment in GBM cell lines causes lowering of HTT protein and RNA. **a** Integration of RNA-seq and proteomics analysis following PRMT5 inhibition in three independent patient-derived GBM lines showing HTT protein and RNA levels are both reduced with PRMT5 inhibitors GSK591 and LY283. Graph represents average fold change (log2) after PRMT5 inhibition combining all three G561, G564 and G583 cell lines and both PRMT5 inhibitors GSK591 and LY-283 against SGC2096 (negative control) (n= 3). Gene density is represented by color intensity (“counts”, log10). **b** qPCR analysis of patient-derived GBM cell lines treated with DMSO, 1 µM SGC2096 (negative control), 1 µM GSK591 (PRMT5 inhibitor) and 1 µM LLY-283 (PRMT5 inhibitor) for 5-7 days, n = 3-5 biological replicates of G523, G411 and G561. PRMT5 inhibitors resulted in a significant lowering of total HTT expression levels relative to DMSO-treated cells. Data are the mean ± SEM of four independent biological replicates each comprising four technical replicates. Statistical significance was calculated using two-way ANOVA with Dunnett’s multiple comparisons test compared to the mean of DMSO-treated cells. ns *P* Value=0.997, \*\*\*\**P* Value<0.0001. **c** Representative western blots of G411G411 and G561 showing significant lowering of HTT protein levels upon treatment with 1 µM GSK591 or 1 µM LLY-283. **d** Western blot quantification is shown as bar graph. HTT protein levels normalized to vinculin and plotted relative to levels observed with DMSO control treatment. Data are the mean ± SEM of four independent biological replicates of G523, G411 and G561. Statistical significance was calculated using two-way ANOVA with Dunnett’s multiple comparisons test compared to the mean of DMSO-treated cells. ns *P* Value=0.921, \*\*\*\**P* Value<0.0001.

### PRMT5 inhibition lowers HTT mRNA and protein levels in HD fibroblasts

Prompted by these findings, we next asked whether this phenomenon can also be observed in cell models of HD. We used h-TERT immortalized control and HD patient-derived fibroblast lines^39^, with *HTT* gene CAG repeat tracts between 40-50, close to the most common clinically observed range. PRMT5 inhibition with 1 µM LLY-283 in these lines resulted in a ∼50% decrease of *HTT* mRNA in comparison to DMSO-treated cells (**Figure 2a**), paralleling what we observed in GBM cells. Western blot analysis showed consistent reduction in HTT protein levels across the series of fibroblasts lines tested, including a homozygous HD line (Q50Q40), homozygous WT line (Q21Q18) and three heterozygous HD lines (Q43Q17, Q57Q17, Q66Q16) (**Figure 2b, 2d**). Similar to our previous observations in GBM cell lines^28^, LLY-283 produced a larger reduction in HTT protein levels than GSK591 (**Figure 2c, d**), consistent with greater potency of LLY-283 over GSK591 for inhibition of PRMT5^28^. Titration of LLY-283 in this cell line showed that significant HTT lowering was evident at 10 nM and peaked at 100 nM (**Figure 2b**), closely following the potency of the compound for its *in*-*cell* target inhibition^40^. Cell viability was not significantly affected at any dose of LLY-283 up to 1 µM, demonstrating that the observed HTT lowering is not an artifact of cell death. To investigate whether the expanded allele of HTT was affected differently to the wild-type, we analysed cell lysates using an allele separation western blotand the anti-polyglutamine antibody, MW1^41,42^. This revealed that the PRMT5 inhibitor treatment led to total HTT lowering, with approximately equal decreased levels of both wild-type and expanded HTT (**Figure 2e**).

**Figure 2.**
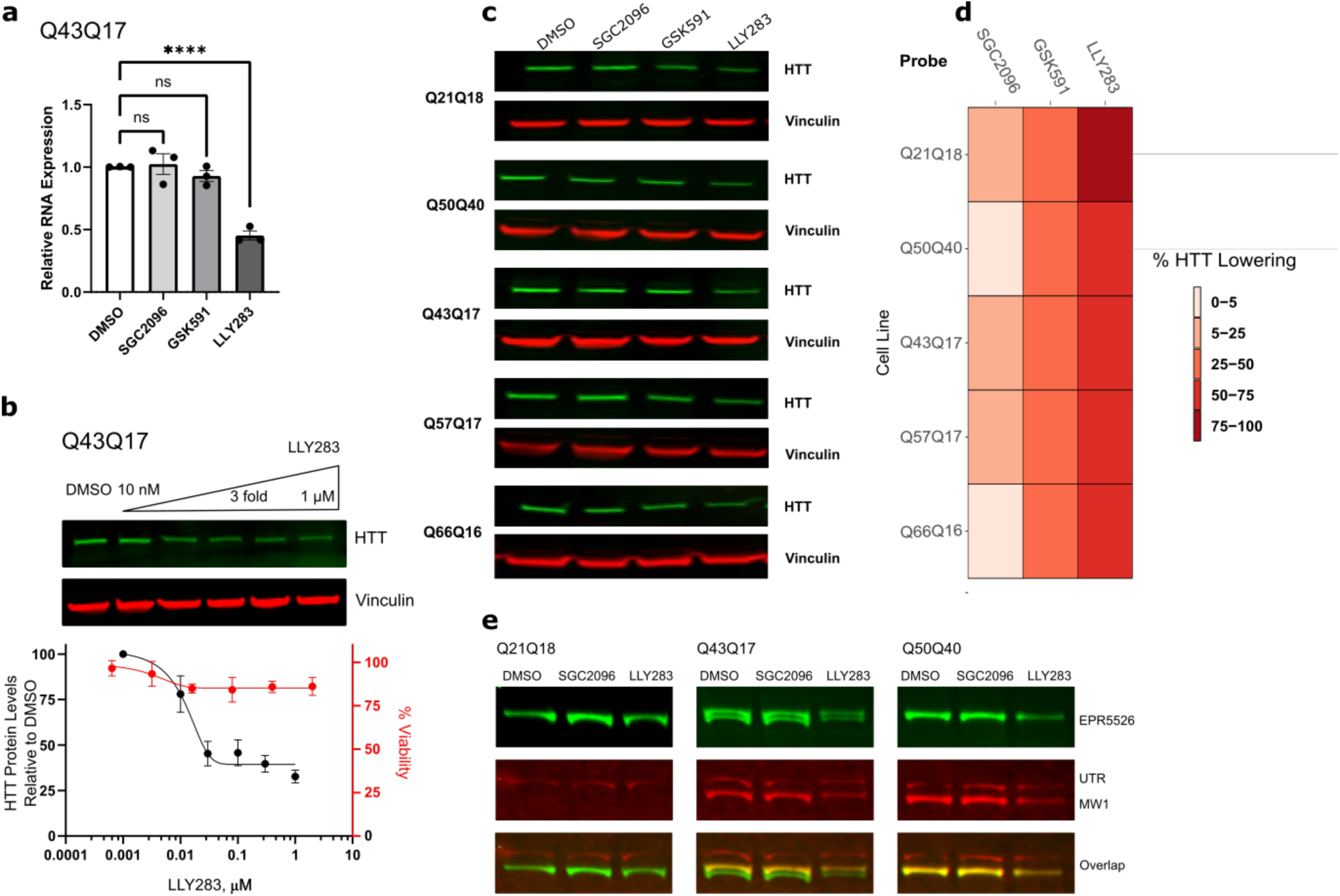
PRMT5 inhibitors lower HTT in control and HD fibroblasts at both the RNA and protein level. **a** Barplot showing the qPCR analysis of TruHD Q43Q17 treated with DMSO, 1 µM SGC2096 (negative control), 1 µM GSK591 (PRMT5 inhibitor), and 1 µM LLY-283 (PRMT5 inhibitor) for 5-7 days. Data are the mean ± SEM of three independent biological replicates each comprising four technical replicates. Statistical significance was calculated using two-way ANOVA with Dunnett’s multiple comparisons test compared to the mean of DMSO-treated cells. ns *P* Value=0.963 and0.474, \*\*\*\**P* Value<0.0001. **b** Top panel: Western blot analysis of HTT and vinculin in control and HD fibroblasts after 5 days of treatment with LLY-283 at different concentrations. Bottom panel: cell viability study of Q43Q17 HD fibroblasts showing no cell death within the concentration range of LLY-283 used in experiments. **c** Representative western blot of HTT and vinculin showing consistent HTT lowering upon treatment with 1 µM LLY-283 in homozygous, (Q50Q40), heterozygous (Q43Q17, Q57Q17 and Q66Q16) and control (Q21Q18) fibroblasts. **d** Heatmap comparing the relative HTT protein lowering in HD (Q50Q40, Q43Q17, Q57Q17 and Q66Q16) and control (Q21Q18) fibroblasts when treated with 1 µM PRMT5 inhibitors (LLY-283 or GSK591) or control. (SGC2096). N= 4-8 biological replicates. **e** Representative western blot probed by EPR5526 for total HTT protein levels, and MW1 for mutant HTT levels in HD fibroblasts. PRMT5 inhibition lowered total HTT levels in HD fibroblasts expressing mutant HTT (Q50Q40, Q43Q17) and control fibroblasts expressing wild-type HTT (Q21Q18). Utrophin (UTRN) was used as a loading control.

### PRMT5 inhibition induces retention of *HTT* introns 9 and 10

Our previous study showed PRMT5 inhibitor treatment induces widespread mis-splicing in GBM cells. Therefore, we reasoned that changes to *HTT* splicing could be the mechanism by which *HTT* mRNA and protein levels are lowered by PRMT5 inhibition. Data mining of our bulk GBM RNA-seq data revealed that two introns (9 and 10) of the *HTT* were mis-spliced upon PRMT5 inhibition, with inclusion levels prominently increased in GBM cells treated with PRMT5 inhibitors (**Figure 3a, Suppl Figure 1**). Remaining introns of the HTT gene analyzed were unaffected (for example, intron 39, **Figure 3b**). Experimental validation of intron retention by qPCR showed dramatic and significant increases for both introns 9 and 10 following PRMT5 inhibition in GBM cells (**Figure 3c, d**) compared to control introns (**Figure 3e**). Visual inspection of the read density across introns 9 and 10 revealed that the intronic reads were diminishing towards the second half of intron 10 (**Figure 3a**). Intriguingly, we found the same trend by analysing RNA-seq data from A549 lung cancer cells treated with GSK591 (**Suppl. Figure 2**) from another study that investigated the effect of PRMT inhibitors on intron retention^27^. Furthermore, there was a visible decrease in exonic reads after exon 10 in both GBM and A549 cells, suggestive of transcription termination (**Suppl. Figure 1** and **Suppl. Figure 2**). This suggests that the PRMT5 inhibitor-induced *HTT* intron retention may lead to transcription termination and mRNA truncation through intronic polyA site (PAS) activation, leading to premature cleavage and polyadenylation (PCPA) of the *HTT* mRNA.

**Figure 3.**
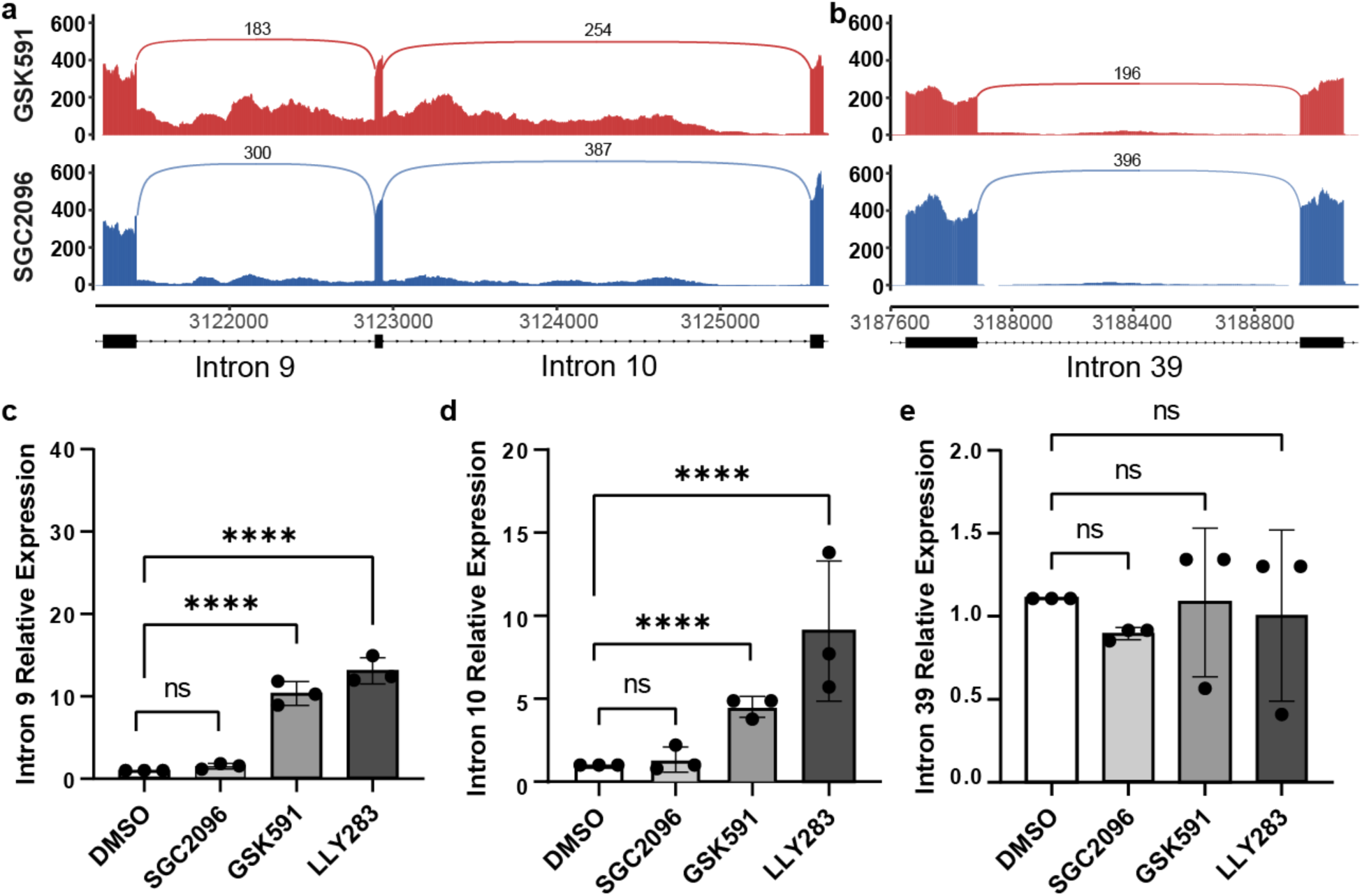
Increase in specific intron retention events in the *HTT* transcript after PRMT5 chemical inhibition. ***a*** Intron retention data in the HTT gene presented as sashimi plots^43^ were extracted from bulk RNA-seq data from patient-derived GBM stem cells treated with 1µM GSK591 or SGC2096 as described previously^28^. **b** *Intron retention* data for *HTT* intron 39 as a control. For both **a** and **b**, X-axis represents the genomic coordinates along the chromosome and Y-axis shows the normalized read count at each position. The lines with numbers connecting the individual exons represent the number of reads that map to the exon-exon junction. **C** Bar plots showing the relative expression of ***c*** *intron 9, **d** intron 10,* and ***e*** *intron 39* as Ct values, (Y-axis) across different conditions (X-axis). Data are the mean ± SEM of three independent biological replicates each comprising four technical replicates. Statistical significance was calculated using two-way ANOVA with Dunnett’s multiple comparisons test compared to the mean of DMSO-treated cells. *P Value ≥ 0.05 is considered insignificant. ****P value < 0.0001*

### PRMT5 inhibition activates multiple cryptic intronic PAS leading to premature cleavage and polyadenylation of *HTT* mRNA

To further explore PCPA as a possible mechanism of intron retention, visual inspection of the sequences of *HTT* introns 9 and 10 revealed six potential alternative polyadenylation (APA) motifs, four clustered toward the distal part of intron 10 (labeled 10A-10D in **Figure 4a**) and two in intron 9 (labeled 9A-9B, **Figure 4e**). We designed qPCR primers upstream of these sites and ran qPCRs using total RNA from GBM stem cell line G523. We used an oligo-dT primer bearing a P7 adapter for the RT step and the P7 adaptor primer coupled to gene-specific primers for the PCR steps as highlighted in **Figure 4b**^44^. We observed very prominent and significant induction of these intronic PAS sites in *HTT* intron 10 upon PRMT5 inhibition compared to the canonical *HTT* PAS sites (**Figure 4c**). Next, we queried the polyA-site database^45^ for the presence of experimentally curated APA sites in the *HTT* gene and found the vast majority mapped within *HTT* introns 9 and 10 (**Table 1**). We then mined single-cell (sc)RNA-seq data derived from GBM stem cells treated with LLY-283 and analyzed PCPA in the *HTT* transcripts. Strikingly, all of the identified major APA events, mapped within *HTT* introns 9 and 10^38^ (**Figure 4d**, **Table 1**). As one of the top-ranked sites by fold change and P-value mapped to intron 9 (labeled 9C in **Figure 4d**, **Table 1**), we designed primers for that site (**Figure 4e**) and performed qPCR using the same RT reaction described above. We saw prominent activation of the intron 9 distal PAS site corresponding to the same site we identified by scRNA-seq (**Figure 4f**, right panel). Furthermore, we found two additional, more proximal, PAS sites in intron 10 (**Figure 4d**, **Table 1**) that we validated by qPCR (**Figure 4f**, left panel). In conclusion, these data collectively point to the induction of PCPA within introns 9 and 10 of *HTT* following PRMT5 inhibition.

**Figure 4.**
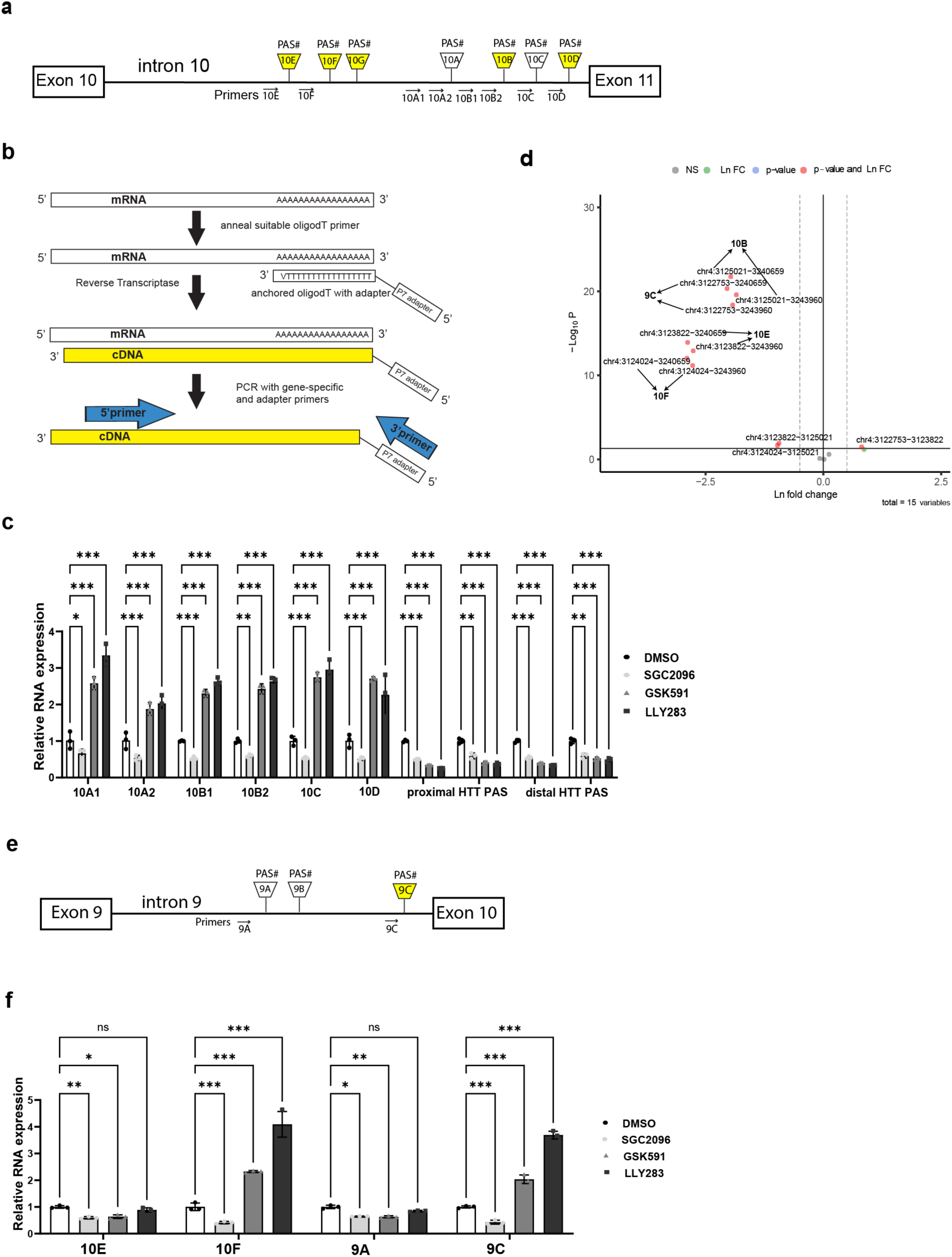
PRMT5 inhibition induces activation of APA sites in HTT introns 9 and 10. **a** Schematic of intron 10 APA sites detected in this study by visual scanning for polyA motifs and our scRNA-seq data analysis. Yellow indicates sites that have been experimentally validated sites. **b** *S*chem*atic o*f *t*h*e RT*-q*P*C*R-*b*ase*d *approa*c*h for* the detection and val*i*dation *o*f 3’ *cD*NA ends. **c** Validation of the potential APA sites using RNA from GBM cells treated with the indicated compounds. For reference, we designed two primers specific for each of the 2 canonical HTT sites (indicated as proximal and distal PAS). Data were analyzed using 2-way ANOVA. Data are shown as mean ± s.d.; n = 3 biological replicates. Asterisks indicate statistical significance. **d** Volcano plot of APA events identified in HTT gene using scRNA-seq data from GBM cells treated with LLY-283 or DMSO for all the APA pairs between proximal and distal Polyadenylation sites. 15 pairwise APA events were observed for the HTT gene. The X-axis denotes the natural logarithm (Ln) fold change of distal to proximal PAs utilization. Hence, the negative values indicate proximal PAs usage and positive values indicate distal PAs usage. We see significant usage of intronic PA sites located on Intron 9 (chr4:3122753) and 10 (chr4:3125021) of the HTT gene for the LLY-283 treated cell lines compared to the controls. **e** Schematic of potential PAS sites mapping in HTT intron 9. Yellow indicates sites that have been experimentally validated. **f** qPCR validation of HTT introns 9 and 10 (additional) PAS sites. Data were analyzed using 2-way ANOVA. Data are shown as mean ± s.d.; N = 3 biological replicates. Asterisks indicate statistical significance based on calculated P-values. *\*P Value = 0.1, **P Value <0.01, and ****P Value<0.0001*.

**Table 1.**
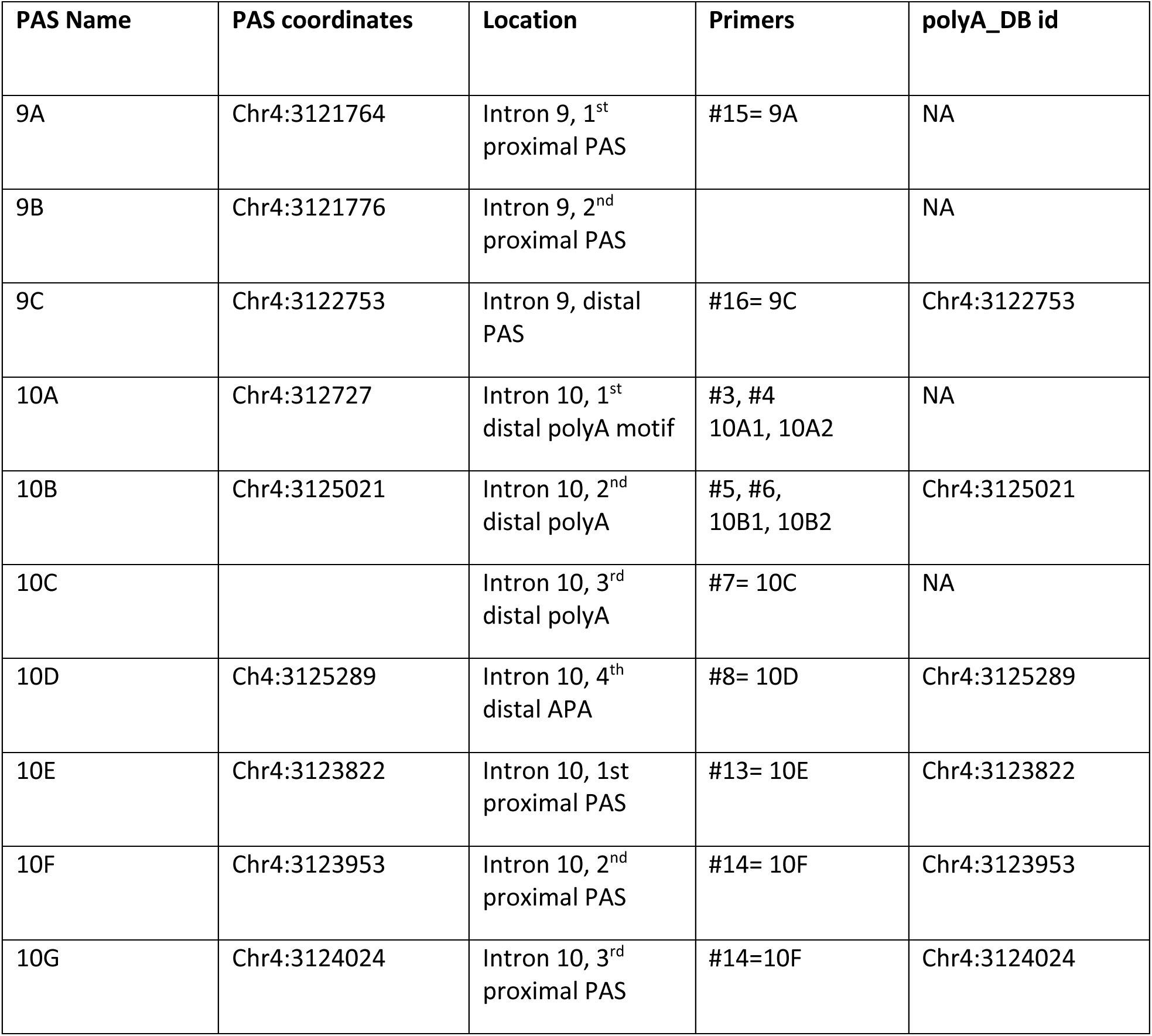
Coordinates of APA sites (PAS) in the *HTT* gene identified by scRNA-seq analysis of two patient-derived GBM lines treated with 1 μM LLY-283 or DMSO.

| PAS Name | PAS coordinates | Location | Primers | polyA_DB id |
| --- | --- | --- | --- | --- |
| 9A | Chr4:3121764 | Intron 9, 1<sup>st</sup><br>proximal PAS | #15= 9A | NA |
| 9B | Chr4:3121776 | Intron 9, 2<sup>nd</sup><br>proximal PAS |  | NA |
| 9C | Chr4:3122753 | Intron 9, distal<br>PAS | #16= 9C | Chr4:3122753 |
| 10A | Chr4:312727 | Intron 10, 1<sup>st</sup><br>distal polyA motif | #3, #4<br>10A1, 10A2 | NA |
| 10B | Chr4:3125021 | Intron 10, 2<sup>nd</sup><br>distal polyA | #5, #6,<br>10B1, 10B2 | Chr4:3125021 |
| 10C |  | Intron 10, 3<sup>rd</sup><br>distal polyA | #7= 10C | NA |
| 10D | Chr4:3125289 | Intron 10, 4<sup>th</sup><br>distal APA | #8= 10D | Chr4:3125289 |
| 10E | Chr4:3123822 | Intron 10, 1st<br>proximal PAS | #13= 10E | Chr4:3123822 |
| 10F | Chr4:3123953 | Intron 10, 2<sup>nd</sup><br>proximal PAS | #14= 10F | Chr4:3123953 |
| 10G | Chr4:3124024 | Intron 10, 3<sup>rd</sup><br>proximal PAS | #14=10F | Chr4:3124024 |

### Increased expression of APA *HTT* isoforms during neuronal differentiation

We were able to detect truncated APA *HTT* isoforms with 3’UTRs composed of intron 9 and 10 sequences in the untreated control conditions in GBM cells, so we reasoned they are likely expressed in specific neuronal tissues. To investigate this possibility, we mined VastDB, an alternative splicing database containing a large compendium of vertebrate gene splice variants and their expression data across different human tissues^46^. Data mining revealed increased levels of *HTT* transcripts with intron 9 and 10 sequences retained during early neuronal development (**Figure 5a**). Strikingly, PRMT5 expression showed the inverse trend i.e. was high in neural stem cells and neuronal progenitor cells (NPCs) but decreased drastically during early embryonic neuronal development (**Figure 5b**). Prompted by this, we mined publicly available neuronal RNA-seq datasets^47–49^ for expression of *HTT* transcripts retaining these introns. We found higher levels of *HTT* transcripts with intron 9 and 10 sequences in differentiated neurons as compared to earlier, less differentiated cell types, represented by human induced pluripotent stem cells (hiPSCs) and neural progenitor cells (NPCs) (**Figure 5c**). Interestingly, mixed neuron/astrocyte populations, which are indicative of inefficient or incomplete neuronal differentiation (Terryn et al, 2020) did not show this increase (**Figure 5c**). Concomitant with this increase in truncated *HTT* transcripts bearing intron 9 and 10 sequences, we saw a significant decrease of PRMT5 expression in neurons compared to cells earlier in the differentiation process (**Figure 5d**). Decreased PRMT5 expression during neuronal differentiation was also observed by Braun *et al*^26^.

We also analyzed a series of isogenic iPSC-derived HD neuronal cell models^50,51^ and found stepwise increases in HTT intron 9 and 10 retention levels during neuronal differentiation from human embryonic stem cells (hESCs) to neurons (**Suppl. Figure 3a**). In the same neuronal differentiation series, we found that the expression of PRMT5 and its cofactor WDR77 decreased during neuronal differentiation (**Suppl**. **Figure 3b**). We conclude that the truncated APA *HTT* transcript isoforms bearing intron 9 and 10 retention are increasingly prevalent through the course of neuronal differentiation and inversely correlate with PRMT5/WDR77 levels.

### PRMT5 inhibition induces neuronal differentiation in GBM stem cells

In order to ascertain the functional impact of PRMT5 downregulation in the differentiation capacity of GBM stem cells we generated a neuronal differentiation reporter assay by using CRISPR-Cas9 genome editing to insert an H2B-CITRINE coding sequence immediately into the TUBB3 locus at the TSS of the human glioblastoma stem cell line G523. 10-14 day treatment with 1μM GSK591 or LLY-283 induced a dramatic induction of the TUBB3-Citrine reporter in these cells (**Figure 5e**). This indicates that PRMT5 activity is required to keep the cells cycling and undifferentiated. Thus, the decrease in PRMT5 expression we observed in neurons (**Figure 5b, d**) is functionally required for neurogenesis and is further correlated to induction of the truncated APA HTT isoform.

**Figure 5.**
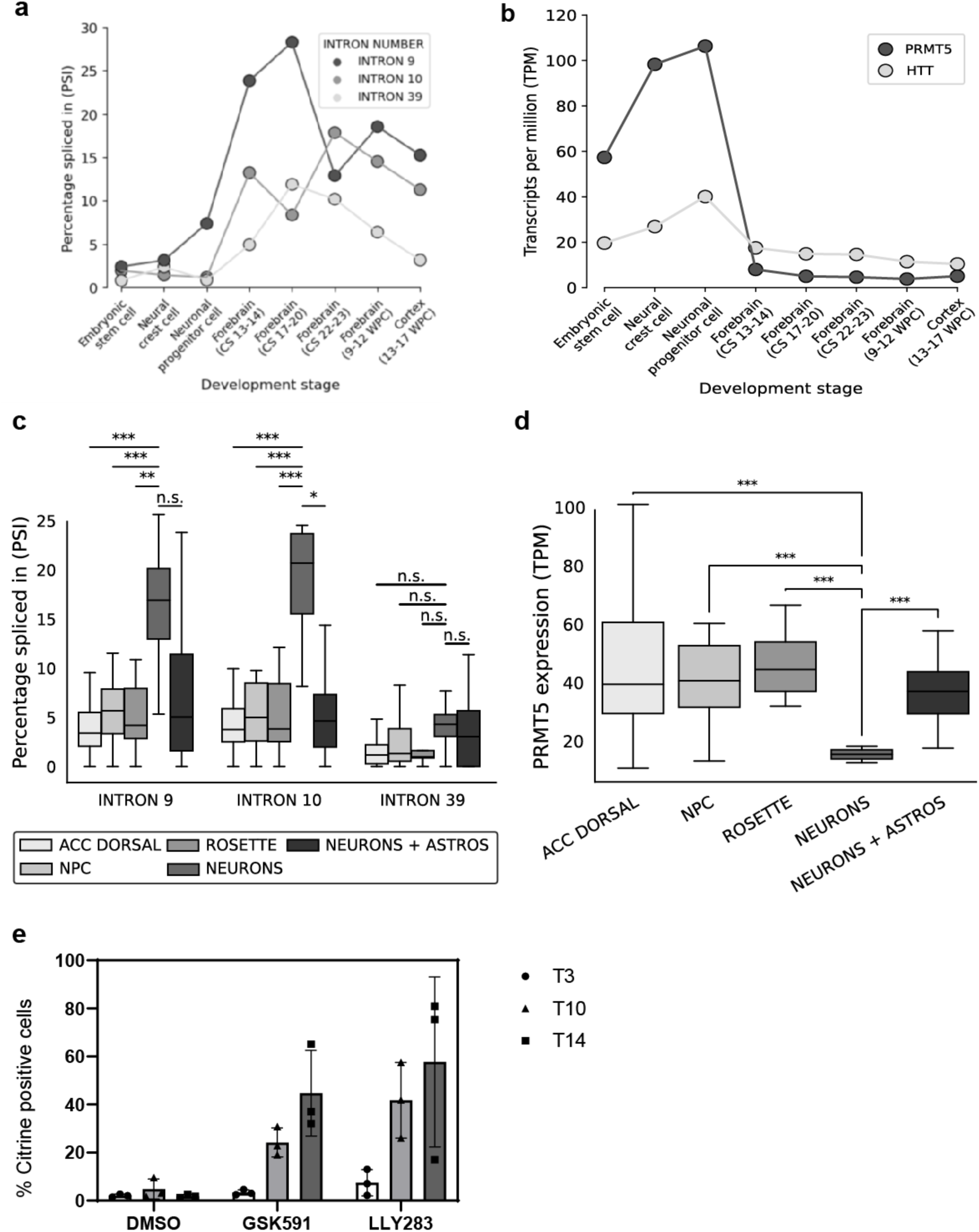
Increased expression of truncated, APA HTT isoforms correlate with lower PRMT5 levels during neuronal differentiation. **a** HTT intron retention events during brain tissue development. Lineplot showing the measured PSI values (Y-axis) for each of the introns of interest (medium and dark gray) at different stages of neuronal development (X-axis). **b** Boxplot showing the percentage spliced-in (PSI, Y-axis) values for each of the evaluated HTT introns (X-axis) across four distinct stages of neuronal development (early differentiating cells represented as ACC dorsal in white, NPC in light grey, rosettes in medium grey, neurons in grey and mixture of neurons plus astrocytes in dark grey). The bars indicate the statistical significance calculated when comparing neurons with other tissues using two-sample independent t-test with unequal variance, with asterisks indicating a *P*-value lower than 0.05 and a ΔPSI (PSI_neuron_ – PSI_target_) higher than 10. **c** Boxplot showing the expression in transcripts per million (TPM, Y-axis) for each of the evaluated introns (X-axis) across the same four different cell types in distinct stages of neuronal development as in panel **b** above. The bars indicate the statistical significance calculated when comparing neurons with other tissues, with asterisks indicating a *P*-value lower than 0.05 and a log2FoldChange(ExpNeuron / ExpTarget) higher than 1 or lower than -1. **d** PRMT5 inhibition in GBM stem cells induces neuronal differentiation. Tubulin B3-Citrin fluorescence reporter assay in GBM stem cell line G523 showing time-dependent induction of neuronal differentiation after treatment with either 1 µM GSK591 or 1 µM LLY283 for the indicated amount of time (T). Cells were collected and processed by flow cytometry at 3, 10 and 14 days of treatment as indicated.

## Discussion

Here, we report the alternative polyadenylation of *HTT* transcripts as a novel downstream process regulated by the arginine methyltransferase, PRMT5. We identify PRMT5 inhibition as a novel way to lower *HTT* mRNA and protein. This was evident for both full-length wild-type *HTT* as well as expanded *HTT* alleles. The effect was mediated by impaired intron splicing leading to PCPA of the *HTT* transcript.

PRMT5 inhibition has been reported to induce widespread mis-splicing in various cell types. A major splicing subtype affected by loss in PRMT5 activity are intron retentions. Intron retentions can be detrimental for gene expression as they not only affect mRNA stability by introducing premature stop codons that lower gene and protein expression by NMD, but can also lead to cryptic cleavage of mRNA and activation of polyadenylation sites, ultimately resulting in mRNA truncation or novel short isoforms^52,53^. Here, we identified several novel intronic cryptic PAS in introns 9 and 10 of the *HTT* gene that are activated by PRMT5 inhibition. This results in PCPA of the *HTT* mRNA which ultimately leads to the decrease in full-length HTT protein levels. This mechanism contrasts with the recently reported splicing inhibitors which lower HTT protein levels by incorporating a novel pseudoexon that introduces premature termination codons, resulting in NMD.

We identify a dramatic activation of a cluster of cryptic intronic PAS sites in introns 9 and 10 of the *HTT* transcript upon PRMT5 inhibition. Although we observed this phenomenon in GBM stem cells and fibroblasts, it is likely conserved in many other cell types. In support of this notion, the majority of intronic *HTT* PAS sites found in the polyA database (a compendium of experimentally validated PAS sites)^45^ map to *HTT* introns 9 and 10 and are identical to the ones found in GBM stem cells and fibroblasts. Moreover, the same HTT introns were identified to be highly retained in reads from a transcriptomic study of HTT splicing isoforms in post-mortem brains. Notably, the reads in introns 9 and 10 in that study showed highly abundant transcripts nearly equal in depth to that of flanking exons^10^. Additional evidence for transcription in introns 9 and 10 has been documented in GENCODE v21^54^. Altogether, this evidence points to the activation of a cluster of cryptic APA sites in *HTT* introns 9 and 10, leading to premature termination and mRNA truncation. This mechanism, termed preT-IR, for premature termination-intron retained mRNAs, was recently described in SFPQ null ALS^53^. Notably, it seems to be prevalent in neuronal mRNAs and the preT-IR transcripts accumulate in neurites^53^.

PRMT5 regulates the assembly of the spliceosome by symmetrically dimethylating SmD1, SmD3 and SmB/B’ proteins, all snRNP components of the spliceosome. The majority of PRMT5 substrates have a role in RNA processing^55^. The spliceosome machinery interacts with RNA polymerase to enhance transcription speed and elongation rate particularly across long introns in long genes^56^. Recently, the U1 spliceosome has been reported to control RNA pol II elongation speed and the ability of RNA pol II to elongate through long AT-rich introns^56^. A propensity for slower transcription and weak splicing may result in the activation and utilization of multiple proximal polyadenylation sites present within these introns. As previously reported, we believe that the weak 5’-splice sites of these introns render them more susceptible to retention after PRMT5 inhibition^25,26^. This very much resembles the U1 telescripting model put forward by Dreyfuss and colleagues^57,58^ observed with U1 snRNP inhibition, where proximal intronic PAS sites are being de-repressed^57,59^. Interestingly, this effect was more evident in long neuronal genes bearing long introns^60^, and PCPA induction was prevalent in activated neurons, stem cells and cancer cells^58^. Importantly, neuronal activation caused a transient U1 snRNP shortage, leading to activation of proximal PCPA and transcript shortening^58^. This model fits perfectly with our findings of a transient PRMT5 decrease during neuronal differentiation accompanied by *HTT* mRNA shortening via proximal APA and PCPA induction. It also suggests that PRMT5 inhibition may result in widespread APA and potential transcriptome shortening via a similar mechanism. In addition, PRMT5 has been reported to directly methylate mRNA cleavage and polyadenylation factors^61^ and RNA polymerase elongation factors such as SPT5 as well as the C-terminal domain (CTD) of RNA polymerase II itself^62^, thus directly impacting transcription termination. Together, our data support a model where PRMT5 inhibition results in de-repression of proximal intronic PAS sites in the *HTT* gene.

We found that the truncated PAS mRNA levels increase during neuronal differentiation, concomitant with a decrease in PRMT5 levels. Previous studies had also found lower PRMT5 levels during neuronal differentiation that were associated with increased levels of intron retention in neurons^26^. Moreover, this decrease in PRMT5 expression is important for neurogenesis as inhibition of PRMT5 activity potently induced the neuronal marker tubulin B3 in GBM stem cells. Thus, lower PRMT5 expression/activity in neurons seems to be a requirement for neuronal differentiation to proceed. This is consistent with previous reports for PRMT5, where higher PRMT5 activity is required in neural stem cells and neural progenitor cells, whereas it decreases in differentiating neurons as well as oligodendrocytes and glial cells^26,68^. The decrease in PRMT5 expression in neurons was accompanied with increased *HTT* intron retention and proximal APA activation, leading to a truncated *HTT* mRNA isoform and lower HTT protein levels. This suggests that lowering full-length HTT via the production of a truncated isoform is a consequence of lower PRMT5 activity during neuronal differentiation. Although this seems counterintuitive at first glance for a protein mutated in neurodegenerative disease, there is evidence showing that HTT is required for neuronal stem cell division and lack of HTT promotes neural stem cell differentiation^30^. Further studies will be required to investigate the role of HTT isoforms during neuronal differentiation. Nevertheless, it is tempting to speculate that the PRMT5-induced isoform switching mechanism we identified here may be a more rapid, malleable and reversible alternative of limiting the production of full-length HTT instead of attempting to directly disrupt transcriptional control, which may require a longer time and expend unnecessary cellular resources. This isoform switching mechanism has the advantage of being tightly linked to transient spliceosomal component imbalances, which may be essential for the transition from neuronal stem cell to neuronal progenitor and subsequently to mature neurons^58^, and possibly during neuronal regeneration^70^. We thus believe our findings provide important insights in HTT regulation during neuronal development and differentiation.

## Materials and Methods

### Cell culture

Immortalised control and HD patient-derived fibroblasts were a kind gift from Professor Ray Truant and have been previously described^39^. TruHD Q43Q17, TruHD Q50Q40, TruHD Q57Q17 and TruHD Q66Q16 were derived from HD patients while TruHD Q21Q17 was derived from a control subject. These cells were cultured in DMEM (Life Technologies #10370) with 15% (v/v) fetal bovine serum (FBS; Life Technologies), 1X GlutaMAX (Life Technologies #35050) and 1x penicillin Streptomycin antibiotics. Cells were grown at 37°C with 5% CO_2_.

G523, G411 and G561 Glioblastoma (GBM) cell lines were grown and maintained as previously described^28,71^. GBM cells were grown in serum-free Neurocult NS-A Basal medium (Stem Cell Technologies #05750) that is supplemented with 2mM L-Glutamine (Wisent #609—65-EL), 75 μg/ml BSA (Life Technologies #15260-037), B27 supplement (Life Technologies #17504-44), rhEGF (10 ng/ml, Stem Cell Technologies #E9644), bFGF (10 ng/mL, Stem Cell Technologies #02634), N2-hormone mix (Thermo Fischer) and 2 µg/ml heparin sulfate (Sigma #H3149). Upon reaching confluence, cells were dissociated via mechanical trituration or with Accutase (Sigma, A6964) and re-plated on 10 cm plates (Corning) pre-coated with poly-L-orthinine (Sigma #P4957, 20 minutes) and laminin (Sigma #L2020).

### Treatment with PRMT5 inhibitors and control compounds

GBM and HD patient fibroblasts were treated with PRMT5 inhibitors in 6 well plates (Costar 6-well Clear TC-treated Multiple Well Plates, Corning) unless stated otherwise. Culture media was refreshed every 2—3 days with ×1 treatment.

HD Patient fibroblasts were plated in 384-well plates (Corning) at a density of 1000 cells/ well, with four technical replicates per dose, and cultured across ten dilution series (ranging from 1 nM to 10 µM) of PRMT5 inhibitors for seven days. The effects of PRMT5 inhibitors on fibroblast cell viability was measured using CellTiter Glo luminescent cell viability assay reagent (Promega, catalogue number: G7570). Cells were grown in 60 µl media. At the endpoint, plates were left to equilibrate to room temperature for 20 minutes, then 25 ul of CellTiter Glo reagent was added to each well and mixed for a further 30 min protected from light. The luminescent signal intensity was subsequently read on a CLARIOstar plate reader (BMG LABTECH). Cell viability values were normalized to that of the DMSO-treated controls for each line. Each data point was plotted using three independent biological replicates.

### Protein extraction and western blot

Lysis buffer containing 20 mM Tris-HCl (pH 8), 0.1% Triton X-100, 150 mM NaCl, 1 mM EDTA, 10 mM MgCl_2_, 1% SDS, protease inhibitors cocktail (in-house), and benzonase (in-house) was used to harvest total protein for western blot. Lysis buffer was added directly to cells on the plate and an aliquot was obtained to determine protein concentration using Pierce™ BCA Protein Assay Kit (Catalog number: 23225). For all western blots, 15–35 µg of protein was loaded on NuPAGE 4–12% Bis-Tris protein mini gels (catalog number NP0321BOX) and transferred onto polyvinylidene difluoride (PVDF) membranes. Primary antibodies used were anti-HTT (Abcam, #EPR5526, 1:1000; CST D7F7 #5656 1:1000), anti-vinculin Antibody 7F9 (Santa Cruz, #sc-73614), and anti-GAPDH (EMD Millipore, MAB374, 1:5000). Secondary antibodies used were IRDye® 680RD anti-mouse IgG (LI-COR, 926-68072, 1:5000) and IRDye®800CW anti-rabbit IgG (LI-COR, 926-32211, 1:5000). Each western blot was replicated in at least three independent experiments. Membranes were visualized on an Odyssey® CLx Imaging System (LI-COR). Full, uncropped images of the western bots are shown in the Source data file.

### Allele separation western blot

To quantify and compare levels of full-length wild-type HTT and mutant HTT, we optimised an allele separating western blot from^41^. 15-35 µg of lysate were run on 3–8% Tris-acetate gradient protein gels (Catalog #: EA03785BOX) for 24h at 120V at 4^0^C until the 250 kDa ladder band run out of the gel. Cold running buffer was refreshed twice during the run (25mMTris base, 190 mM glycine, 3.5 mM SDS and 10.7vmM β-mercaptoethanol). Protein was transferred onto 0.2 um PVDF membrane (BioRad) using the NuPAGE transfer buffer (25 mM Bicine, 25 mM Bis-Tris, 1 mM EDTA, and 15% methanol) at 70V for 2h. Membrane was then blocked for 1h with 5% skimmed milk in phosphate-buffered saline with 0.05% Tween-20 (PBST) at room temperature. Primary antibodies used were for mutant HTT MW1 (Milipore # MABN2427). Secondary antibody use and membrane visualization are the same as described above.

### RNA extraction and Quantitative real-time PCR

CDNA was generated using iScript gDNA Clear cDNA Synthesis kit (Bio-Rad) according to the manufacturer’s protocol. 500 ng-1 µg of total RNA was treated initially with DNAse to eliminate any residual genomic DNA contamination and then incubated with iScript reverse transcription supermix to produce cDNA required for the subsequent steps. For all qRT-PCR reactions, a no RT control was included as per the manufacturer’s recommendations to control for residual genomic DNA contamination in reactions.

A 5× dilution was then made by adding 80 µl of double-distilled water, and 2 µl of synthesized cDNA was used per reaction. During qRT–PCR, cDNA template and LUNA master mix (NEB) were added into white PCR reaction tubes and placed in a CFX Maestro (Bio-Rad). All experiments were performed in at least four technical replicates and data was obtained from 3-4 independent experiments. Primers were designed to measure fully spliced transcripts and intron-containing transcripts, and all primer sequences are provided in **Supplementary table 2**. For data analysis, the threshold cycle (Ct) values obtained during qRT–PCR was used to calculate the ΔCtΔCt of intron-containing transcripts relative to the geometric mean of the expression of housekeeping genes GAPDH and U6. Relative RNA expression (ΔΔCt) of intron-containing was calculated by comparing PRMT5 inhibitor treated versus DMSO-treated samples. Statistical comparison was calculated using two-way ANOVA with Dunnett’s multiple comparisons test compared to the mean of DMSO-treated cells. Data were collected on a Bio-Rad CFX96 touch and analyzed by Bio-Rad CFX Manager (v3.1.1517.0823) and GraphPad prism 8.

### Proteomics analysis

Mass-spec analysis was carried out as described previously^28^. ^28^. Briefly, GSC lines G561, G564 and G583 were treated for 6 days with either 1 μM of GSK591, 1μM LLY-283 or the inactive control, SGC2096 (n=3). Proteins were then extracted with denaturing urea buffer in the presence of protease and phosphatase inhibitors and digested with trypsin. Desalted peptides were analyzed by mass spectrometry (MS, “1D separation”) on the Orbitrap Fusion instrument using liquid chromatography (LC) gradients. Peptide identification and label-free quantification were done in PEAKS (version X) considering relative abundance of all identified peptides for each protein. Subsequent data analysis was done in R (version 4.2.3). The mass spectrometry proteomics data have been deposited to the ProteomeXchange Consortium (http://proteomecentral.proteomexchange.org) and is accessible via the PRIDE partner repository with the dataset identifier PXD021635^28^.

### Splicing data analysis

Intron retention data in the HTT gene presented as sashimi plots were extracted from bulk RNA-seq data from patient-derived GBM stem cells treated with 1uM GSK591 or SGC2096 as previously described^28^.

We analyzed 297 published samples in total as follows:

139 samples from hiPSC^47^, 40 samples from hESC^72^, 83 samples from hPSC-derived, 35 samples from hiPSC^48^, and 288 samples from hiPSC.

Each dataset was evaluated to identify intron retention events (measured in PSI, or percentage spliced in) and gene expression values (in TPM, or transcripts per million). We utilized the vast-tools suite to compute intron retention events and gene expression values for each sample. Subsequently, we compared the PSI levels of introns 9 and 10, as well as the gene expression levels, across various cell types and stages of differentiation to identify statistically significant alterations between these variables. Furthermore, we conducted a thorough analysis of the VASTdb tissue/cell-line database to determine reliable intron retention patterns for the specific events in a panel that accurately depicts different stages of neuronal development. We encountered a notable constraint in the quantity of reads collected in the exon-exon and/or exon-intron junctions of the HTT gene in certain instances. To address this, we utilized the “vast-tools combine” module to merge replicates and samples from identical experimental settings to perform adequate statistical inferences. The post-processing data analysis and figure generating stages were performed using custom Python 3.7 scripts, which can be obtained upon request.

The RNA-sequencing of the isoHD allelic differentiation series from Ooi et al^50^ and Tano et al^51^ was used to visualize the expression levels of PRMT5 and WDR77 as well as the introns of HTT. This set included embryonic stem cells (ESCs, n=9), neural progenitor cells (NPCs, n=9) and neurons (NEU, n=12). Briefly, gene level differential expression was performed using DESeq2^74^. Junction level differences in intron 9-10 in HTT were tested using JunctionSeq^75^ normalised to other detected junctions, including introns 59-60 and 64-65. To visualize, the gene and exon/intron counts from all samples were normalized using the variance stabilising transform from DESeq2^74^ and plotted with the boxplots from ggplot2. Boxplots represent the median (line), the 25^th^ and 75^th^ percentiles (hinges) and 1.5 times the inter-quartile range (whiskers). As in Tano et al, the ESCs and NPCs were generated and sequenced as one experiment and are directly comparable. The NEUs were generated and sequenced as a second experiment.

### APA analysis in PRMT5 inhibitor-treated GBM cells

We performed single cell RNA-seq using the 10xGenomics technology in two patient-derived GBM cell lines G549 and G561 treated with vehicle DMSO or 1uM LLY-283. Similar to the workflow in [McKeever et al, 2023] we used <u>MAAPER</u> software package, to assign sequencing reads to the known, curated polyadenylation sites^76^. Then, we used the APAlog^77^software package in pair-wise mode to quantify the APA events. This enabled us to assess APA usage across all known polyadenylation (PA) sites within transcripts. Using APAlog, we calculated the base 2 logarithm (Log_2_) of the fold change to evaluate the significance of differential usage between distal and proximal PA sites.

#### TUBB3 neuronal differentiation reporter assay

To generate a reporter for induction of neuronal lineage differentiation we used CRISPR-Cas9 genome editing to insert an H2B-CITRINE coding sequence immediately into the TUBB3 locus at the TSS of the human glioblastoma stem cell line G523. G523-TUBB3-CITRINE cells were seeded in poly-l-ornithine + laminin coated dishes and treated with either 1 µM GSK591, 1 µM LLY283 or 0.1% DMSO for 14 days with regular media changes. At 3, 10 and 14 days of treatment, cells were collected and fixed in 1% paraformaldehyde. Flow cytometry on fixed cells was performed using either a CytoFlex S or CytoFlex LS system (Beckman Coulter) and analyzed using CytExpert version 2.6 software (Beckman Coulter).

## Author Contributions

M.A.A.: investigation, methodology, analysis, writing; F.E.C.: analysis, methodology, writing; J.C.H.: investigation, methodology, analysis; M.I.M. investigation, methodology, analysis, writing; M.Y.: investigation, methodology, analysis; A.M.S. analysis, methodology; G.M. investigation, methodology, resources, analysis; M.A. investigation, resources, methodology; G.B. investigation, resources, methodology; V.T.: analysis, methodology; S.H.L.: analysis, methodology, supervision; M.S.-O.: analysis, methodology; P.S. investigation, resources, methodology; M.K.: investigation, methodology, resources; L.R.: investigation, methodology, analysis; C.F.B.: investigation, methodology; M.A.P.: supervision, resources; T.P.: supervision, resources, funding acquisition; M.T.: supervision, funding acquisition; S.A.: supervision, resources, funding acquisition; P.B.D. supervision, resources, funding acquisition; G.D.B.: supervision, funding acquisition; K.B.M. supervision, writing; D.B.-L.: supervision, methodology; D.S. supervision, funding acquisition; R.J.H. supervision, methodology, writing; C.H.A.: supervision, funding acquisition, writing; P.P.: supervision, methodology, analysis, writing.

## Acknowledgments

We would like to thank Ray Truant, Rebecca Kurtz, Paul Boutz and Christian Braun for helpful discussions and sharing data. We thank Magdalena Szewczyk and Shili Duan for help with Westerns and tissue culture experiments respectively and Jean-Philippe Brosseau for help with QPCR primer design. M.A.A. was funded by the Princess Margaret Postdoctoral Fellowship and is currently funded by a MITACS Accelerate fellowship. This work was supported by SU2C Canada Cancer Stem Cell Dream Team Research Funding (SU2C-AACR-DT-19-15) provided by the Government of Canada through Genome Canada and the Canadian Institute of Health Research (142434), with supplemental support from the Ontario Institute for Cancer Research through funding provided by the Government of Ontario. Stand Up To Cancer Canada is a program of the Entertainment Industry Foundation Canada. Research funding is administered by the American Association for Cancer Research International-Canada, the scientific partner of SU2C Canada. D.S. was supported by the following grants: NIH/NIGMS R01GM108646 and the Irma T. Hirschl Trust. The Structural Genomics Consortium is a registered charity (no: 1097737) that receives funds from Bayer AG, Boehringer Ingelheim, Bristol Myers Squibb, Genentech, Genome Canada through Ontario Genomics Institute [OGI-196], EU/EFPIA/OICR/McGill/KTH/Diamond Innovative Medicines Initiative 2 Joint Undertaking [EUbOPEN grant 875510], Janssen, Merck KGaA (aka EMD in Canada and US), Pfizer and Takeda.

