## Supplementary Figures 1-3 and Supplementary Table 1 for "PRMT5 is required for full-length *HTT* expression by repressing multiple proximal intronic polyadenylation sites"

Supplementary Figure 1

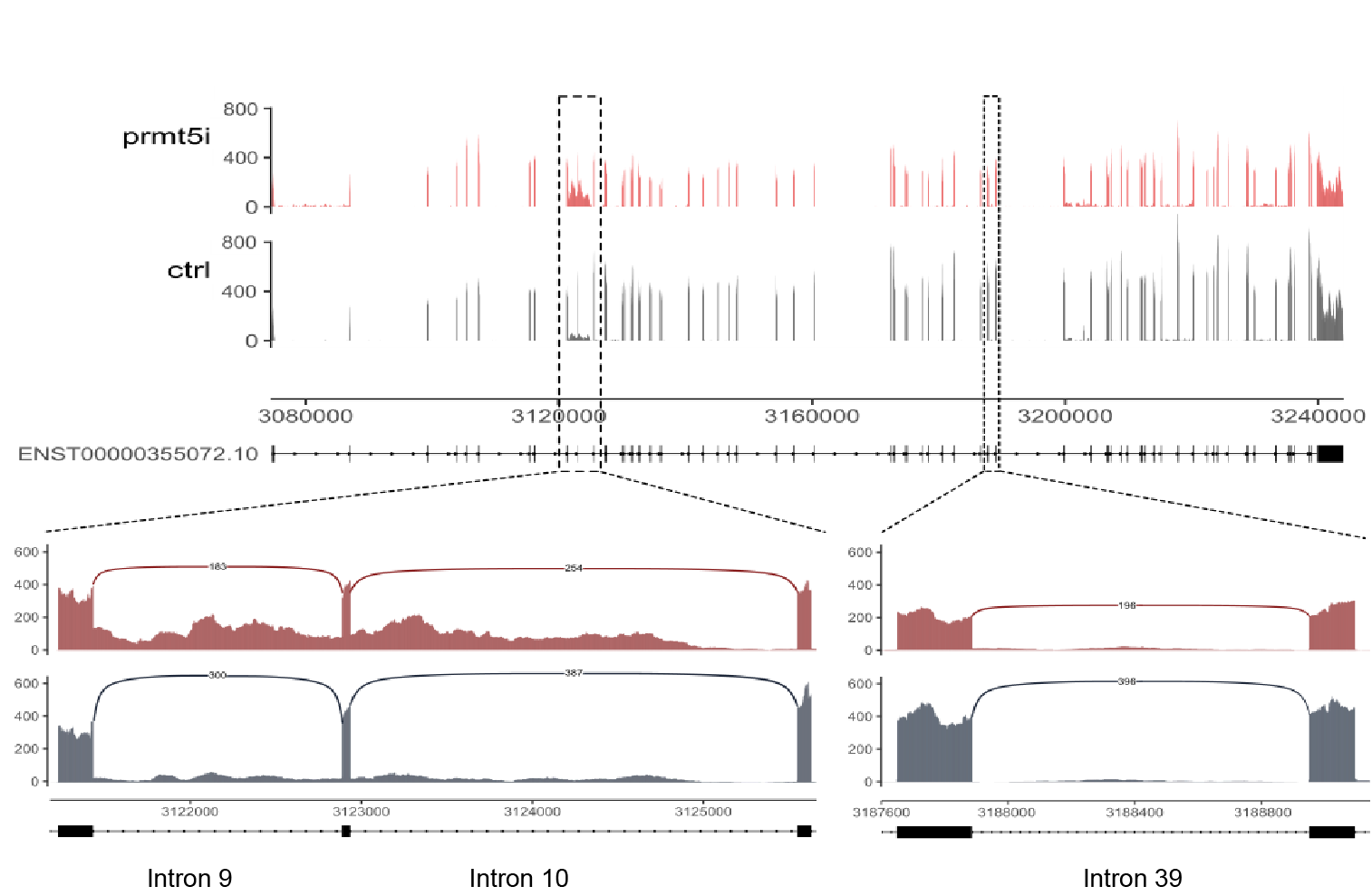

**Supplementary Figure 1**: PRMT5 inhibition results in HTT intron retentions in GBM stem cells

Bedgraph representation of the estimated expression distribution across the HTT gene structure derived from RNA-seq data from GBM cells treated with GSK591 (denoted in red) or SGC2096 (in black), represented by its most abundant isoform (ENST00000355072). The X-axis represents the gene positions along the chromosome and the Y-axis shows the normalized read count at each position. The boxes represent exons and the introns are shown as lines. The zoomed in panels in the bottom left represent introns 9 and 10 whereas intron 39 is on the bottom right.

Supplementary Figure 2:

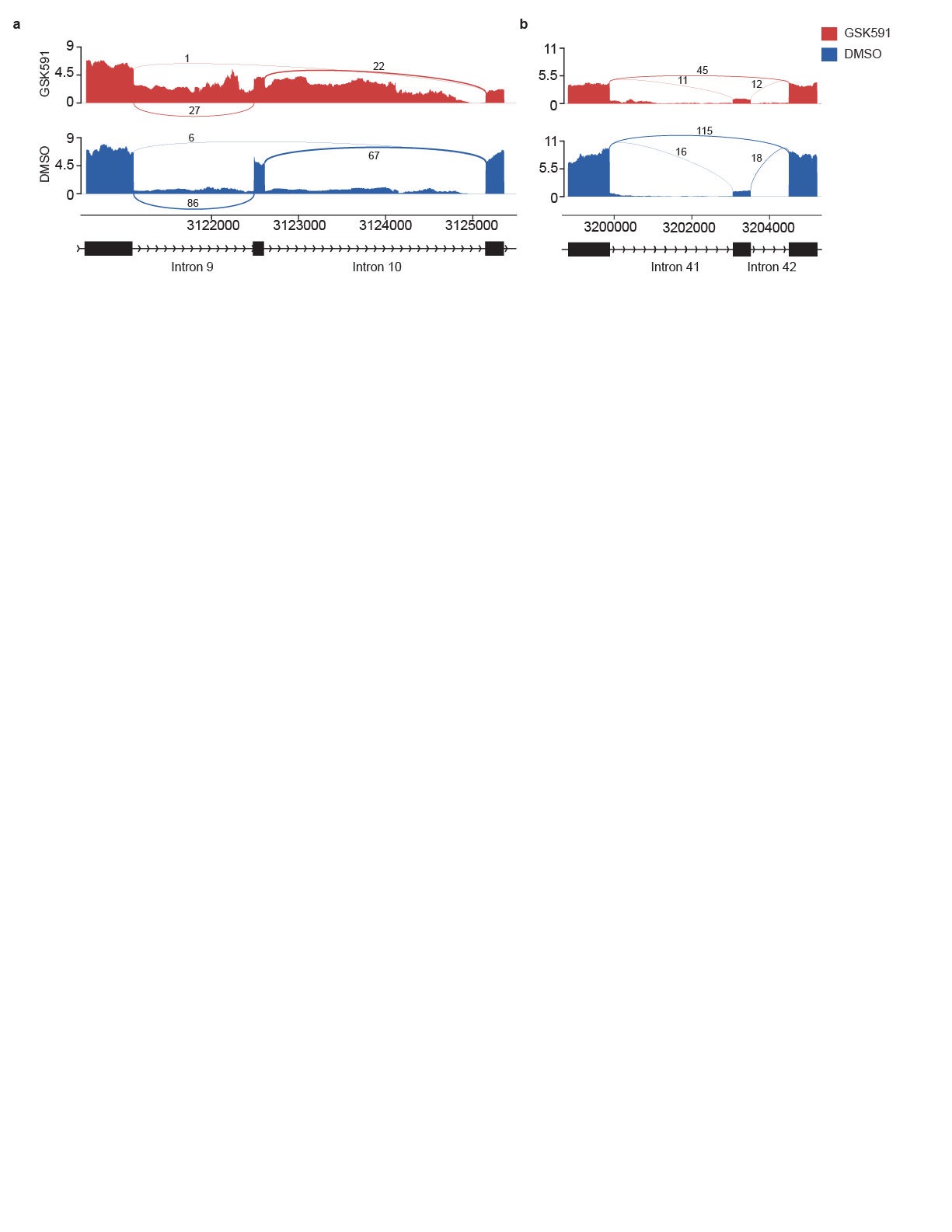

**Supplementary Figure 2. PRMT5 inhibition induces HTT intron retentions in A549 cells**

Sashimi plot representation of the reads distribution across HTT intron 9 and 10 derived from RNA-seq data from A549 cells treated with GSK591 (denoted in red) or DMSO vehicle control (denoted in blue).

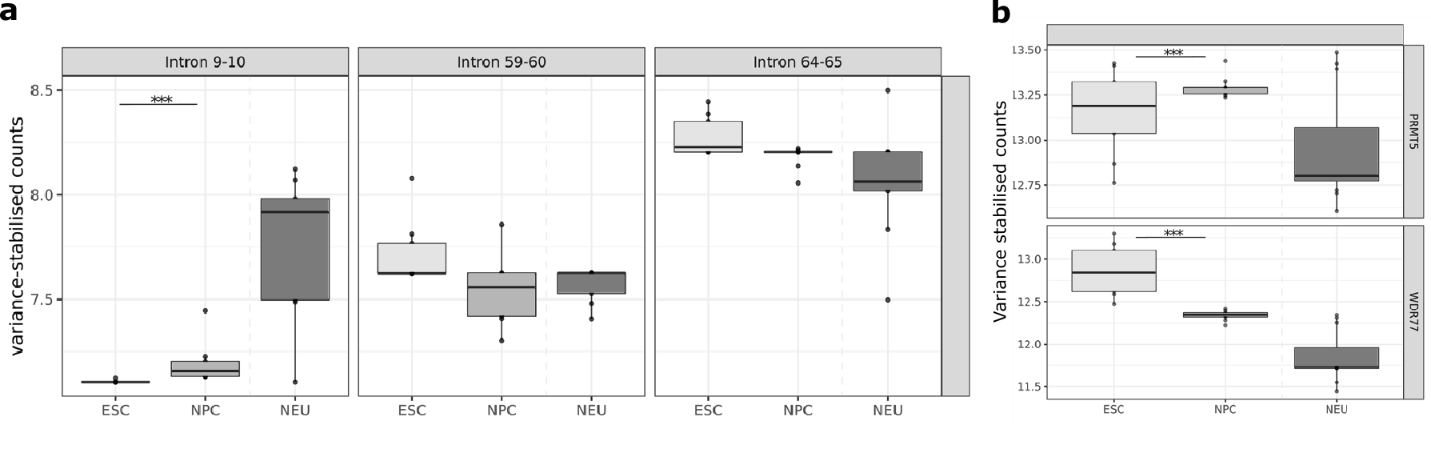

**Supplementary Figure 3. Increased retention of *HTT* intron 9 and 10 sequences during neuronal differentiation of iPSCs.**

**a** Box plots of the normalized gene expression for HTT introns (9-10, 59-60, 64-65) in the ESCs, NPCs, and NEUs from the isoHD allelic series neuronal differentiation in Tano *et al* 2023^51^. The ESCs and NPCs were sequenced together as one experiment and are directly comparable while the NEUs were a separate experiment. **b** Boxplots of the normalized gene expression for PRMT5 and WDR77 from the same isoHD series as in d. Boxplots represent the median (line), the 25th and 75th percentiles (hinges) and 1.5 times the inter-quartile range (whiskers).

**Supplementary Tables**

**Supplementary Table 1**. Splice site scores for annotated HTT splice variants in the VASTDB database.

| VASTDB ID Event | Type | Length | Coordinates | Exon/  Intron # | PSI Average | PSI Range | 5'ss Score | 3'ss Score |
| --- | --- | --- | --- | --- | --- | --- | --- | --- |
| HsaEX0030684 | Exon Skipping | 341 | chr4:3127264-3127604 | Exon 12 | 99.8 | 3.27 | 6.43 | 8.54 |
| HsaEX0030685 | Exon Skipping | 124 | chr4:3129924-3130047 | Exon 13 | 99.49 | 26.17 | 5.32 | 7.43 |
| HsaEX0030686 | Exon Skipping | 143 | chr4:3102606-3102748 | Exon 4 | 3.70 | 100.00 | 4.22 | 6.04 |
| HsaEX0030687 | Exon Skipping | 64 | chr4:3135904-3135967 | Exon 20 | 99.72 | 25.74 | 5.86 | 7.79 |
| HsaEX0030704 | Exon Skipping | 90 | chr4:3202977-3203066 | Exon 42 | 13.91 | 31.10 | 3.77 | 9.10 |
| HsaEX1017274 | Exon Skipping | 146 | chr4:3172580-3172725 | Exon 30 | 0.24 | 26.42 | 11.11 | 6.82 |
| HsaEX1017276 | Exon Skipping | 61 | chr4:3112992-3113052 | Exon 6 | 1.09 | 25.00 | 4.85 | 9.80 |
| HsaEX6083330 | Exon Skipping | 137 | chr4:3180515-3180651 | Exon 36 | 99.72 | 34.50 | 10.51 | 10.77 |
| HsaINT0078263 | Intron Retention | 257 | chr4:3130048-3130304 | Intron 13 | 10.63 | 63.28 | 7.43 | 7.01 |
| HsaINT0078288 | Intron Retention | 1702 | chr4:3180652-3182353 | Intron 36 | 6.72 | 64.06 | 10.77 | 9.08 |
| HsaINT0078291 | Intron Retention | 1064 | chr4:3187887-3188950 | Intron 39 | 8.36 | 61.06 | 9.55 | 8.91 |
| HsaINT0078294 | Intron Retention | 4067 | chr4:3199940-3204006 | Intron 41 | 8.75 | 35.64 | 6.22 | 7.35 |
| HsaINT0078302 | Intron Retention | 1248 | chr4:3212710-3213957 | Intron 49 | 9.74 | 63.30 | 10.06 | 7.02 |
| HsaINT0078305 | Intron Retention | 2553 | chr4:3215212-3217764 | Intron 51 | 7.58 | 47.86 | 9.35 | 8.26 |
| HsaINT0078306 | Intron Retention | 2229 | chr4:3217953-3220181 | intron 52 | 7.16 | 50.65 | 10.10 | 11.49 |
| HsaINT0078312 | Intron Retention | 134 | chr4:3228746-3228879 | Intron 59 | 11.90 | 86.49 | -0.25 | 8.51 |
| HsaINT0078324 | Intron Retention | 1456 | chr4:3121433-3122888 | intron 9 | 8.44 | 87.87 | 1.91 | 9.77 |
| HsaINT0078260 | Intron Retention | 2612 | chr4:3122889-3125629 | Intron 10 | 6.87 | 31.50 | 5.88 | 3.77 |

**Supplemental Table 2**: Primer sequences used for qRT-PCR.

| **Gene** | **Primer name** | **Sequence** |
| --- | --- | --- |
|  | oligodT with P7 adapter for RT | CAAGCAGAAGACGGCATACGAGATTTTTTTTTTTTTTTTTTTTTTTTTVN |
|  | P7 primer | CAAGCAGAAGACGGCATACGAGAT |
| HTT | APA forward primer #3 | GCCTAGCTCTTGTGCCTTT |
| HTT | APA forward primer #4 | TGTCCTGGAGCCTCTCTT |
| HTT | APA forward primer #5 | CACATGTGATGTAGCCACAAAG |
| HTT | APA forward primer #6 | TGAAGAGCAGAAATTAGAAATTCCC |
| HTT | APA forward primer #7 | CCTTCCAGATCATATAATGCTTAAGTTC |
| HTT | APA forward primer #8 | GGCCAATAGCATTGGATCTTTAC |
| HTT | APA forward primer #9 | GCTGTGATGGGCGAGTC |
| HTT | APA forward primer #10 | CAGCAGGGCTGTGATGG |
| HTT | APA forward primer #11 | CGTGTCTGGATGCACAGAT |
| HTT | APA forward primer #12 | ACACTGGCCTGGGTCTC |
| HTT | APA forward primer #13 | TTTGAGCCCAGGAGTTTGAG |
| HTT | APA forward primer #14 | TCCAGTGTGGGTTTGCATAG |
| HTT | APA forward primer #15 | AGGCTGAGGTGGTAGGATT |
| HTT | APA forward primer #16 | GGAATGCGTGCCTTACAAATA |
| HTT | Global F1 | CCCCAAAAGCCATCAGCGAGG |
| HTT | Global R1 | GACTCCACCATTTCTGCCACCA |
| HTT | Intron 9 Forward | TGGGAGTATTGTGGAACTTATAGGCA |
| HTT | Intron 9 Reverse | GCAGGCCCATAATCAATAGAGTGTAA |
| HTT | Intron 10 Forward | CCTGTCCTTTCAAGAAAACAAAAAGGTG |
| HTT | Intron 10 Reverse | CAGTAAGATTCAACACAAGACTCTGATTTC |
| HTT | Intron 39 forward | TTTTTGTTTCTTGGAAG/GTTTC |
| HTT | Intron 39 reverse | GATGTGGATCAGACACATTAGCA |
| PRMT5 | Global F1 | TCCAAGCTGTACAATGAGGTCC |
| PRMT5 | Global R1 | GAGTCTCTGGACGGATACTCAG |
| WDR77 | Global F1 | AGCAGAAGCCTTCGTTGG |
| WDR77 | Global R1 | ATGGAACGGCAGTTGGAG |
| U6 | F | GAGAAGATTAGCATGGCCC |
| U6 | R | AATATGGAACGCTTCACGA |
| GAPDH | F | CATGAGAAGTATGACAACAGCCT |
| GAPDH | R | AGTCCTTCCACGATACCAAAGT |
